# A patient-designed tissue-engineered model of the infiltrative glioblastoma microenvironment

**DOI:** 10.1101/2020.10.02.322735

**Authors:** R. C. Cornelison, J. X. Yuan, K. M. Tate, A. Petrosky, G. F. Beeghly, M. Bloomfield, S. C. Schwager, A. L. Berr, D. Cimini, F. F. Bafakih, J. W. Mandell, B. W. Purow, B. J. Horton, J. M. Munson

**Affiliations:** Department of Biomedical Engineering, Virginia Tech, Blacksburg, VA 24061; Department of Biomedical Engineering, University of Massachusetts Amherst, Amherst, MA 01003; Department of Biomedical Engineering, University of Virginia, Charlottesville, VA 22904; Department of Biological Sciences and Fralin Life Sciences Institute, Virginia Tech, Blacksburg, VA 24061; University of Virginia School of Medicine, Charlottesville, VA, 22903; Department of Pathology, University of Virginia, Charlottesville, VA 22903; Department of Neurology, University of Virginia, Charlottesville, VA 22903; Department of Public Health Sciences, University of Virginia, Charlottesville, VA 22903

## Abstract

Glioblastoma is an aggressive brain cancer characterized by diffuse infiltration. Infiltrated glioma cells persist in the brain post-resection where they interact with glial cells and experience interstitial fluid flow. We recreate this infiltrative microenvironment *in vitro* based on resected patient tumors and examine malignancy metrics (invasion, proliferation, and stemness) in the context of cellular and biophysical factors and therapies. Our 3D tissue-engineered model comprises patient-derived glioma stem cells, human astrocytes and microglia, and interstitial fluid flow. We found flow contributes to all outcomes across seven patient-derived lines, and glial effects are driven by CCL2 and differential glial activation. We conducted a six-drug screen using four outcomes and find expression of putative stemness marker CD71, opposed to viability IC_50_, significantly predicts murine xenograft survival. Our results dispute the paradigm of viability as predictive of drug efficacy. We posit this patient-centric, infiltrative tumor model is a novel advance towards translational personalized medicine.

## Introduction

Glioblastoma (GBM) is the most common and malignant form of primary brain cancer, and clinical treatments have advanced slowly over the last 25 years ago. Introduction of the Stupp protocol (surgical resection, radiation, and oral temozolomide chemotherapy) established the current median GBM patient survival of 15 months (*1*). The difficulty in treating GBM is attributed to diffuse invasion into the surrounding tissue where tumor cells acquire therapy resistance or increased malignancy in response to therapy (*2–4*). Identifying drugs to target and kill these invaded cells has proven challenging, particularly because drug screens often use tumor cells alone on tissue culture plastic – a poor representation of the tumor or invaded brain. Multicellular spheroid cultures may recreate tumor geometry but can overlook elements like stromal cells, space for diffuse tumor spread, and biophysical factors found in the tissue.

The tissue surrounding a tumor, known as the tumor microenvironment (TME), contains cellular and extracellular factors that contribute to cancer progression (*5,6*). In GBM and other cancers, the cellular TME can enrich cancer stem cell populations and increase tumor cell survival, proliferation, invasion, and drug resistance (*7*). Additionally, we and others have shown that the biophysical force known as interstitial fluid flow, which increases during tumorigenesis, stimulates tumor cell invasion (*8–11*). The brain TME is particularly unique because the primary matrix component is hyaluronan as opposed to fibrillary collagen found in carcinomas, and it contains cells unique to the central nervous system like astrocytes and microglia. Unfortunately, these attributes are difficult to recreate in experimental model systems, and orthotopic xenografts are the primary way to study these TME elements in combination. However, these animal models are quite expensive, offer little control over experimental variables, and ultimately may not capture patient heterogeneity or drug response as well as expected (*12*).

An ideal tumor model would be cheaper and offer modular control over experimental variables for elucidating distinct and emergent contributions of the cellular and biophysical TME in GBM progression and therapy. Tissue-engineered models of cancer offer substantial control and potential for high throughput screening, are easily tunable, and are cost-effective compared to animal models. Unlike traditional 2D cell culture, three-dimensional culture systems enable approximating *in vivo* tissue physiology using relevant parenchymal cells, extracellular matrices, and relevant mechanical cues/forces. While there are models containing multiple parenchymal cells reported for breast, ovarian, and pancreatic cancer (*13–15*), most GBM models focus on combining tumor cells with only one other cell population (*16,17*). A recent study reported co-culture of astrocytes, microglia, and tumor cells in a 2D format (*18*), but the architecture of the microenvironment is critical for recreating cellular states found in human GBM (*19*). Furthermore, current models were also developed using arbitrary ratios of tumor cells to other cells, and our recent work established a need to use real cellular ratios since the composition of invasive brain tissue in patients predicts survival (*20*).

Here, we report the rational design of a 3D *in vitro* model of the human GBM TME incorporating human astrocytes, microglia, patient-derived glioblastoma stem cells, and interstitial fluid flow. The cellular ratios are defined from invasive margins of patient resection samples, and the interstitial flow rate is based on previous measurements in small animals (*21*). Our model uses a hyaluronan-based matrix mimicking the primary extracellular matrix of the brain and a tissue culture insert format to enable application of fluid flow and drug therapies at a physiologically-relevant flow rate. This format also enables us to examine invasion, cell death, and phenotypic markers for cancer stem cells, cell proliferation, and glial cell activation, collectively providing a holistic assessment of how the TME influences GBM malignancy. Specifically, we use this model to examine 1) individual and synergistic effects of the cellular and biophysical GBM microenvironment on glioblastoma stem cell outcomes, 2) drug response *in vitro* and prediction of murine xenograft survival *in vivo*, and 3) the relationship between patient-specific glial cell activation and patient cell line phenotypes.

## Results

### Invasive GBM regions primarily contain astrocytes, microglia, and tumor cells

We obtained and analyzed resected tissue samples from 63 patients who underwent treatment for GBM at the University of Virginia Cancer Center from 2010-2015. A neuropathologist determined 40 of the 63 samples to contain sufficiently large regions of tumor-adjacent ‘reactive areas’ (**Figure 1B**). We used hematoxylin and eosin as well as Movat pentachrome staining to assess general tissue properties and chromogenic staining to identify neuroglial cell types (**Figure 1C-F**). We quantified the cells as a percent of total cell fraction. While the patient samples are highly variable (**Figure 1G**), we find both astrocytes (ALDH1L1+) and microglia (Iba1+) constitute approximately 18-19% of tumor-adjacent, reactive areas (**Figure 1D**). We previously showed area fractions of these cell types in adjacent regions significantly correlated with patient survival (*20*), but number fractions do not correlate (**Figure S1**). In sequential sections, we determine approximately 75% of the reactive areas are neurons (∼1%), oligodendrocytes (∼16%), and otherwise unidentified cells (∼58%, based on mucin coverage). To build the ‘average’ TME model, we set astrocytes and microglia at 1:1 and a final glioma:astrocyte:microglia ratio at 75:12.5:12.5 or 6:1:1. A cartoon of the general tissue model design is shown in **Figure 1A**.

**Figure 1.**
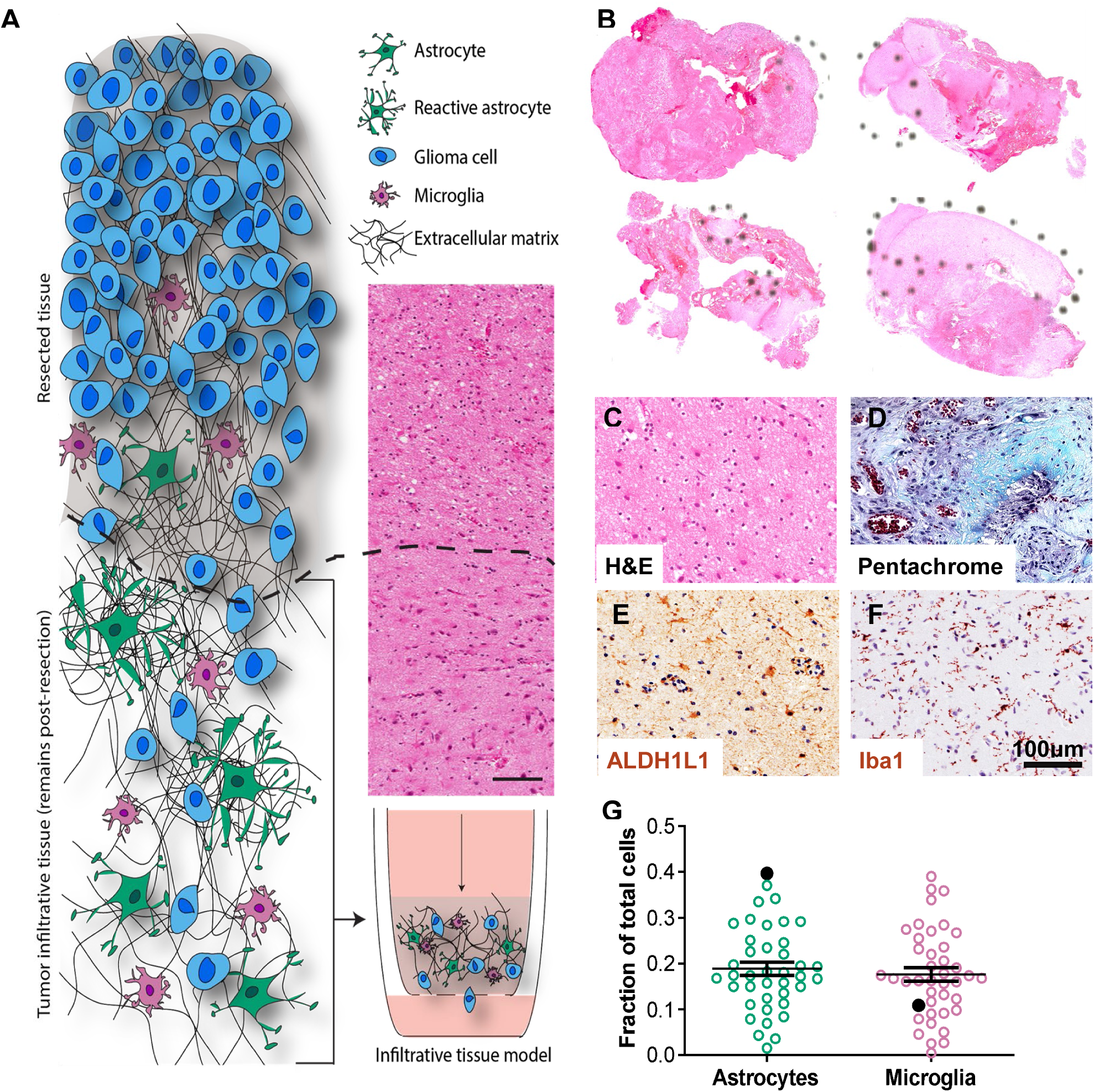
Histological quantification of the invasive human glioblastoma microenvironment for in vitro model development. A) Cartoon of the invasive tumor border and the patient-driven approach to develop a relevant model. B) Representative bright field scans of patient resection samples stained with hematoxylin and eosin (H&E), with dashed circles showing tumor-adjacent regions identified by a neuropathologist. C-F) Representative bright field images of chromogenic stains on serial patient samples for H&E (C), movat pentachrome (matrix and mucin staining, D), ALDH1L1 (astrocytes, E), and Iba1 (microglia, F). G) Cell number quantification from our patient cohort samples (N=40) for astrocytes and microglia, represented as fraction of total nuclei count. Solid black circles show data from a select patient.

### Tissue culture insert model recreates in vivo xenograft phenotypes

Based on our histological findings in patient GBM samples, we developed a tissue culture insert model of the human GBM TME using tri-culture of patient-derived glioblastoma stem cells (GSCs), human primary astrocytes and immortalized microglia in a hyaluronan-based hydrogel. This model incorporates interstitial flow to mimic physiological drug delivery (by applying a pressure head to the transwell) and is compatible with multiparametric flow cytometry analysis of tumor cell metrics like proliferation (Ki67+), stemness (CD71+), and cell death (**Figure 2A**). Tumor cell invasion is also quantifiable by imaging the underside of the tissue culture insert, and the gels can be stained for immunocytochemical analysis (**Figure 2B**).

**Figure 2.**
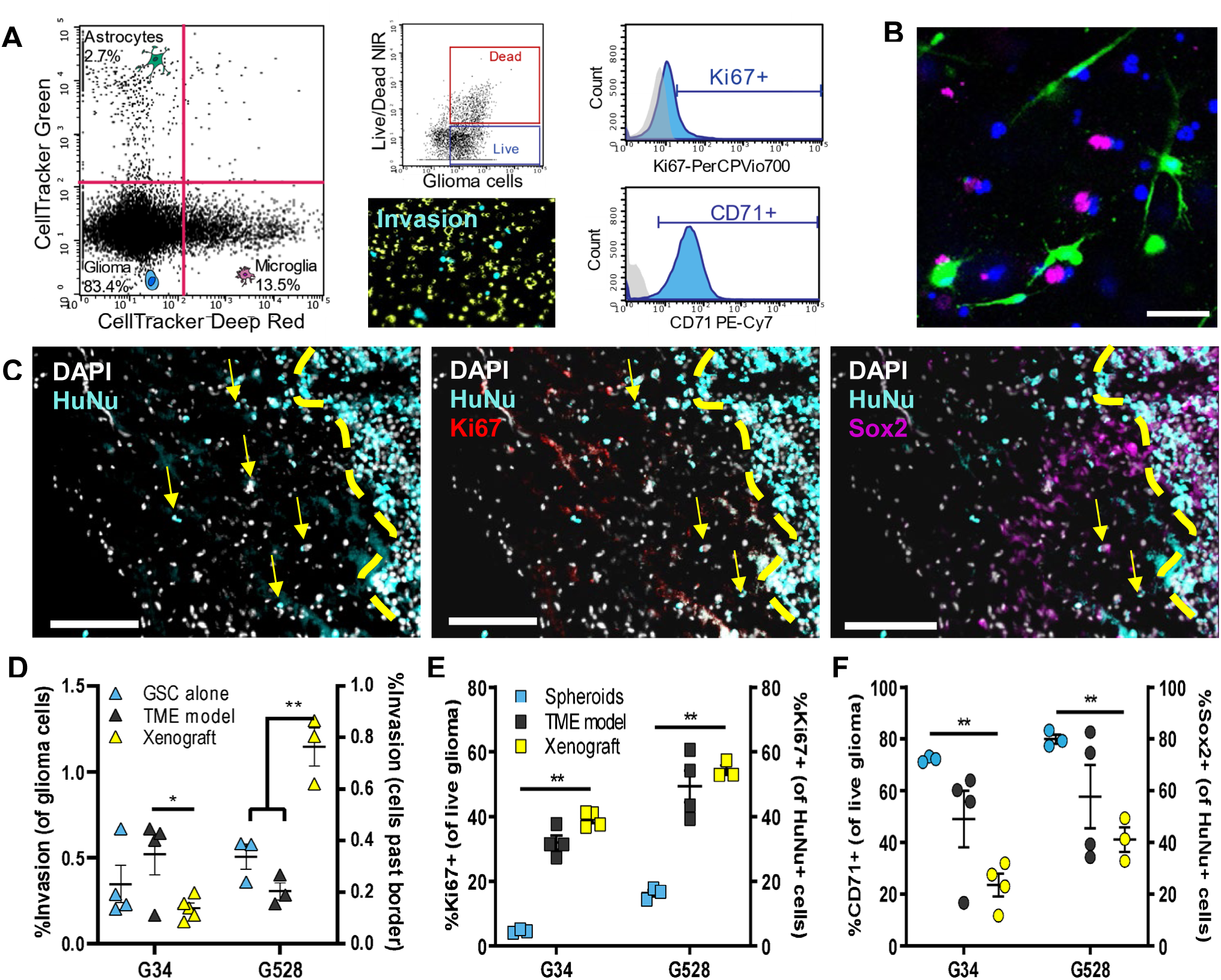
Tunable model of the human invasive TME enables multiplexed analysis of glioma markers and more closely recreates *in vivo* expression than spheroid culture. A) Representative flow cytometry plots showing the ability to distinguish between astrocyte, microglia, and glioma cell populations and determine glioma-specific proliferation (Ki67^+^), stemness (CD71^+^), and cell death. Also shown is a representative fluorescence image of the underside of the porous tissue culture insert used to quantify invasion. B) Representative fluorescence image within the gel showing the three distinct cell populations with glioma (blue), astrocytes (green), and microglia (magenta). Scale bar is 50 μm. C) Representative images of an orthotopic xenograft tissue sample used for immunohistochemical staining and counting of invaded cancer cells beyond the tumor border (based on human nuclear antigen, HuNu^+^). Sections were also stained for Ki67 and Sox2 to quantify the percent of proliferating and stem-like cells of those invaded cancer cells. Scale bars are 200 μm. D-F) Comparison of the %invasion (D), %Ki67^+^ cells (E), and %CD71^+^ cells (F) for three cancer models. For D, invasion was quantified from GSCs encapsulated alone in the gel above a transwell membrane. For E-F, we compare GSCs cultured as spheroids vs. incorporated into the TME model vs. from *in vivo* tissue sections. Keys for E-F are the same. All comparisons to *in vivo* sections are conducted using unpaired t-tests with *p<0.05 and **p<0.01 for n=4.

We first optimize the fluorescent labeling, co-culture, and re-isolation of GSCs with astrocytes and microglia (**Figure S2**). Next, we show the validity of this model by comparing tumor cell expression of proliferative and stem cell markers in the model to that in spheroid culture and orthotopic xenografts (**Figure 2C-F**). Cells in spheroid culture are significantly less proliferative but significantly more stem-like than cells in the TME model or in xenografts. The TME (tri-culture) model does not effectively recreate the invasiveness of glioma cells *in vivo*, and invasion may be better assessed *in vitro* using GSC hydrogel monoculture (**Figure 2D**). However, the tri-culture model does approximate the percentage of proliferating and stem-like cells better than spheroid culture for two representative GSC lines (G34 and G528) (**Figure 2D-F**). These data show a higher-complexity TME model better recapitulates certain *in vivo* tumor cell characteristics than spheroid or mono-culture.

### Glioma cell invasion is highly patient-specific and dependent on TME context

We sought to determine the impact of each of our microenvironmental components on the three outcomes. Because infiltrative and drug resistant cells in glioblastoma may be slower-cycling and more stem-like (*22*), we also wanted to determine if invasion, proliferation, and stemness are correlated across our patient lines. In total, we used seven different GSC lines, with characteristics of these lines are shown in **Table S1**. We tested the effect of GSC line (G2, G34, G44, G62, G262, G267, or G528), transport condition (static or flow), and glia (no glia, astrocytes, microglia, or both) on our three outcomes (proliferation, invasion, and stemness) and employed quantile regression modeling of these covariates which accounted for the skewness of our outcomes (**Figure S3; Table S2)**. GSC line is the only covariate involved in significantly contributing to all three outcomes, either alone or interacting with the other covariates, indicating that inter-patient differences are the greatest contributor to invasion, proliferation, and stemness, regardless of other conditions within the model. GSC line (=83.97, p<0.0001), as well as GSC line interactions with glia (*X*^2^=38.59, p=0.0032) or glia and transport, are significant predictors of percent invasion (*X*^2^=29.16, p=0.0464). The lines G2, G34, and G528 consistently show invasion 10-fold higher than the other lines. Flow generally increases invasion when glioma cells are cultured alone, but the responses are more variable once glia are present (**Figure 3A-B**). For example, we see increased invasion of G2 under flow when cultured alone but decreases in invasion under flow when microglia and/or astrocytes are present (**Figure 3C**). With other cells, such as G34, the effects of flow and glia are more summative. Using the model, it is also possible to tune the glial cell ratio to recreate individual patient data, and changes in the ratio of microglia has more impact on glioma cell invasion and stemness than changing the astrocyte ratio (**Figure S4**).

**Figure 3.**
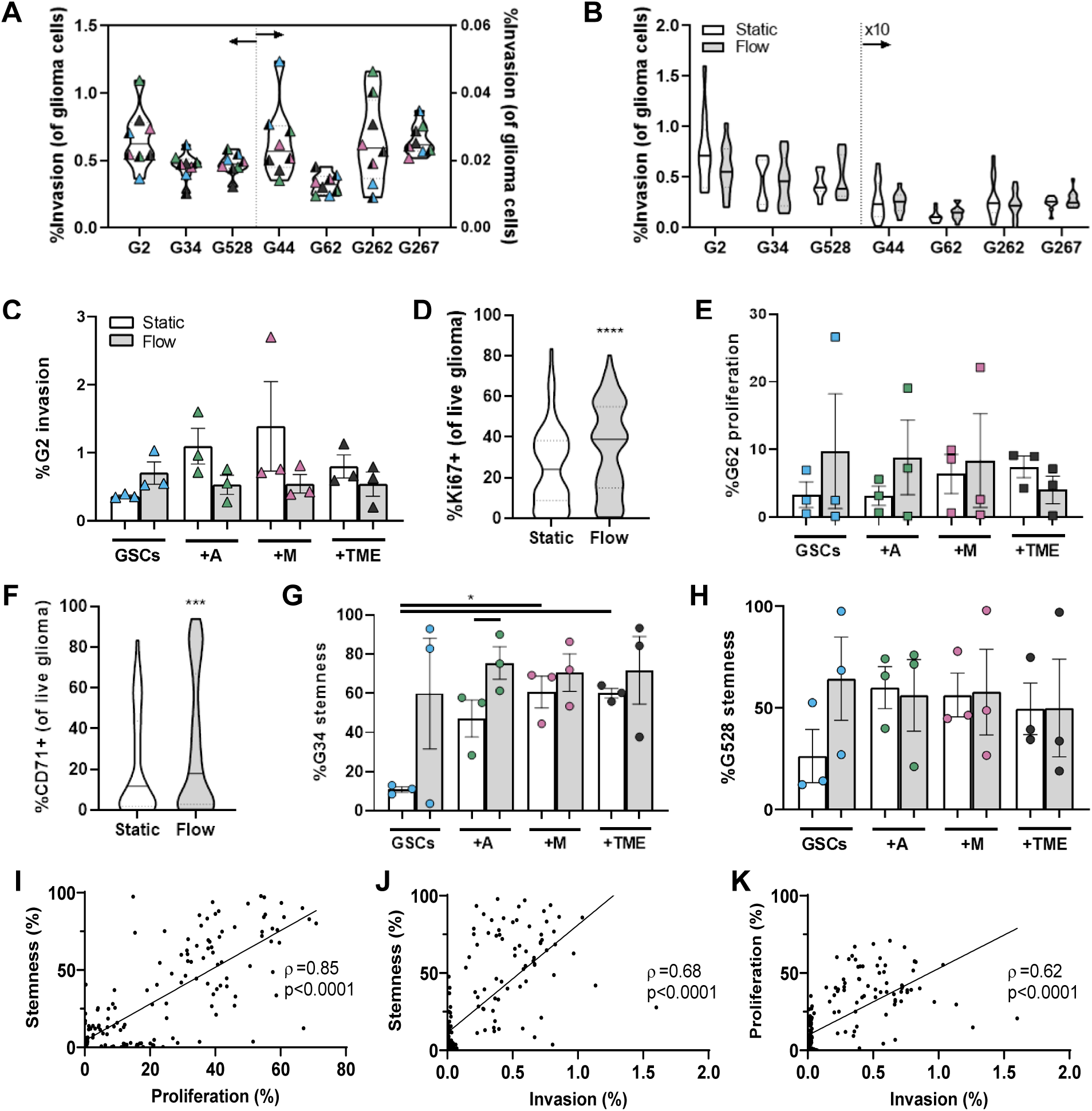
Interstitial flow induces the largest effect on glioma metrics. A) Violin plot of individual invasion data for each cell line in response to TME elements: GSCs alone (blue), +astrocytes (green), +microglia (pink), or +both (gray) in static (solid) vs in flow (half filled). Each point is n=3 technical replicates. Data for G44, G62, G262, and G267 are on a different y-axis scale (right). B) Violin plots showing collective invasion data for all conditions (+astrocytes, microglia, or both) in static (white) vs in flow (gray). Data for G44, G62, G262, and G267 are multiplied by a factor of 10 to be on the same scale. C) Plot of G2 invasion with each TME element to show what is clustered in graphs A and B. D) Percent of Ki67^+^ proliferating cells for all GSC lines in response to TME elements in static (white) vs in flow (gray). E) Plot of G62 proliferation with each TME element to show what is clustered in D. F) Percent of CD71^+^ stem cells for all GSC lines in response to TME elements in static (white) vs in flow (gray). G-H) Specific plots of G34 and G528 stemness from data within (F). Above statistics performed by paired t-tests with *p<0.05, ***p<0.001 and ****p<0.0001. I) Correlation plot of stemness (CD71^+^) and proliferation (Ki67^+^). J) Correlation plot of stemness (CD71^+^) and invasion. K) Correlation plot of proliferation (Ki67^+^) and invasion. Pearson’s coefficients are shown in respective plots.

### Proliferation response is sensitive to cell line or flow

Proliferation of tumor cells is a major factor in the survival of patients, and we find percent proliferation is highly dependent on the patient GSC line (*X*^2^=36.22, p<0.0001) or patient GSC as it interacts with transport (*X*^2^=29.81, p<0.0001). Proliferation increases under flow compared to static conditions for two lines, G2 (t=3.47, p<0.001) and G34 (t=3.5, p<0.001), but significantly decreases under flow for G44 (t=-2.45, p<0.001) and G267 (t=-2.28, p<0.05). Collectively, interstitial flow significantly increases glioma cell proliferation across all lines (**Figure 3D; Figure S3; Table S2**). While glia do not significantly contribute to glioma cell proliferation across all lines, they do within single lines. For example, G62 shows increased proliferation with the addition of either astrocytes or microglia under flow, but significantly decreased proliferation with both glia present under flow (t=-1.99, p<0.05) (**Figure 3E**).

### Stem-like populations respond to each element of the TME model

The presence of cancer stem cells can be an important indicator of tumor growth and recurrence (*23*). We find expression of the stem-like marker CD71 is significantly modeled by the interaction of GSC line with transport condition (X^2^=64.68, p<0.0001), glia components (X^2^=86.62, p<0.0001), and all three covariates (X^2^=40.26, p<0.0019). There are also statistically significant interactions between transport condition and either glia or GSC line. Considering all lines together, the addition of flow and/or glial cells increases stem populations in GSC compared to the static control (**Figure 3F**). These changes vary in effect size and direction across the cell lines (**Figure 3G, H**), suggesting that expression of the stemness marker CD71 is a more sensitive metric compared to invasion and proliferation for response to each of the TME variables tested (astrocytes, microglia, and flow). To understand how these outcomes interrelate, we conducted correlation analyses for the outcomes across all conditions. Putative stemness and proliferation displayed a very strong positive correlation with each other (**Figure 3I**, Spearmon R=0.85, p<0.0001). Stemness also displayed a positive correlation with invasion (**Figure 3J**, R=0.68, p<0.0001), as did proliferation (**Figure 3K**, R=0.62, p<0.0001), but with a moderate effect. Importantly, these effects are all positively correlated, indicating that ‘malignancy’ metrics increase similarly regardless of the experimental parameters.

### Pro-tumorigenic effects of the glioma TME are driven by CCL2

The TME is known to cross-communicate with glioma cells and drive tumor progression (*24*). To examine glioma-glia communication in our model, we analyzed the cellular secretomes in monoculture versus tri-culture. Based on a 44-plex cytokine array (Luminex), we find glial cells are a major source of pro-tumorigenic CCL2 while glioma cells express CXCL1 and CXCL8. These cytokines are upregulated to varying degrees in tri-culture, dependent upon the glioma line (G2, G34, or G528) (**Figure 4A**). Glial co-culture with G2 or G34 induces upregulation of all three cytokines, while G528 showed only minor effects except for a decrease in CCL2. Results in the literature suggest each cytokine can contribute to glioma cell invasion and stemness, but the effects on proliferation are less documented (**Figure 4B**).

**Figure 4.**
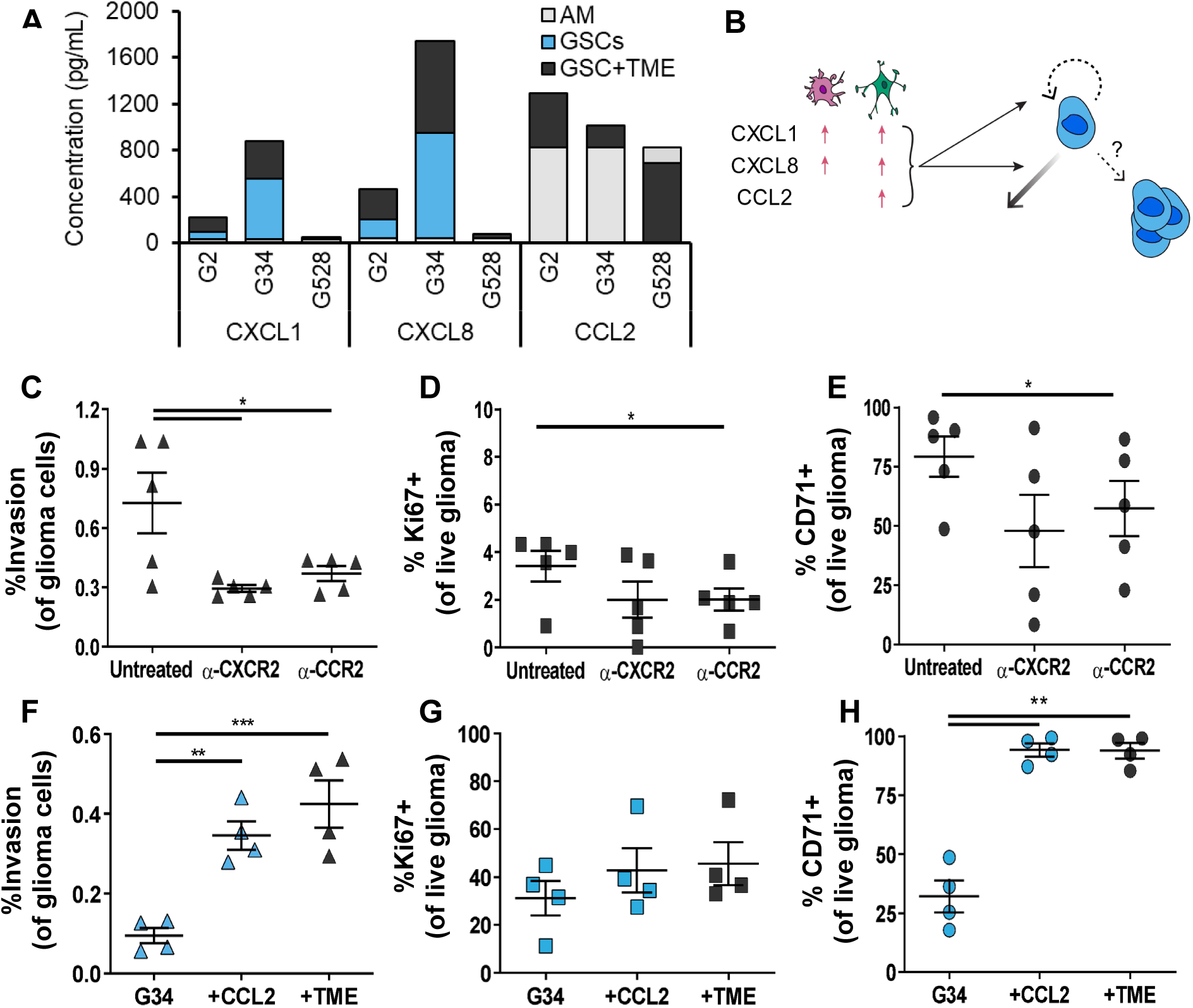
Cytokine expression is altered in tri-culture, with CCL2 able to recreate many of the TME effects on glioma outcomes. A) Cytokine expression based on Luminex array for astrocytes and microglia alone (AM) and in combination with GSC lines G2, G34, or G528. The baseline for each G34 monoculture is subtracted from the respective tri-culture condition to show synergistic opposed to additive effects. B) Schematic showing what is known about the three cytokines pertaining to glial expression following activation and the known effects on glioma invasion, proliferation and stemness (counterclockwise from left). C-E) Effects of adding blocking antibodies against the cytokine receptors CXCR2 (for CXCL1 and CXCL8) and CCR2 (for CCL2) in tri-culture outcomes of invasion (C), proliferation (D), and stemness (E). F-H) Effects of adding CCL2 to GSC monoculture hydrogels vs. the full cellular TME for invasion (F), proliferation (G), and stemness (H). Statistics performed by paired t-tests with *p<0.05, **p<0.01, and ***p<0.001.

Blocking each cytokine directly using antibodies shows varied results (**Figure S5**), potentially because cytokines diffuse faster than antibodies and can be sequestered by the matrix; therefore, we used antibodies against the relevant receptors instead (CXCR2 for CXCL1/CXCL8 and CCR2 for CCL2). Blocking either CXCR2 or CCR2 decreased invasion of G34, with α-CXCR2 having a significant effect (**Figure 4C**). Conversely, only blockade of CCR2 induced significant decreases in both G34 proliferation and stemness (**Figures 4D, E**). Furthermore, the addition of CCL2 to GSCs in gel alone significantly increases invasion and stemness of G34 without influencing proliferation, which recreates the effects of adding the cellular TME (**Figure 4F-H**). This one cytokine is therefore sufficient to replicate the effects of the TME on glioma metrics.

### Cancer-associated glial activation is patient line-specific

Most research into the TME focuses on how the stromal cells affect the cancer, but, since cancer:stromal crosstalk affects is bidirectional and possibly cyclic, it may be equally important to determine how the cancer affects stromal cells. We used the TME model to test the effects of GSCs on glial cells, focusing on glial activation. We used immunocytochemistry to evaluate expression of GFAP in activated astrocytes and CD68 in activated microglia (**Figure 5A**). The baseline activation in glia-only cultures was low for astrocytes and moderate for microglia, possibly because the microglia are immortalized. The presence of GSCs influenced astrocyte activation the most, with every line significantly increasing the number of GFAP^+^ astrocytes except G2 (**Figure 5B**). The percentage CD68^+^ microglia tended to increase with G528, G2, and G34 but decrease with G262 and G44 (**Figures 5C**). Unexpectedly, astrocyte activation strongly negatively correlated with microglia activation across all cell lines (**Figure 5D**). There was also an interesting trend for the response to cluster by the original tumor subtype, but more data is necessary for definitive conclusions.

**Figure 5.**
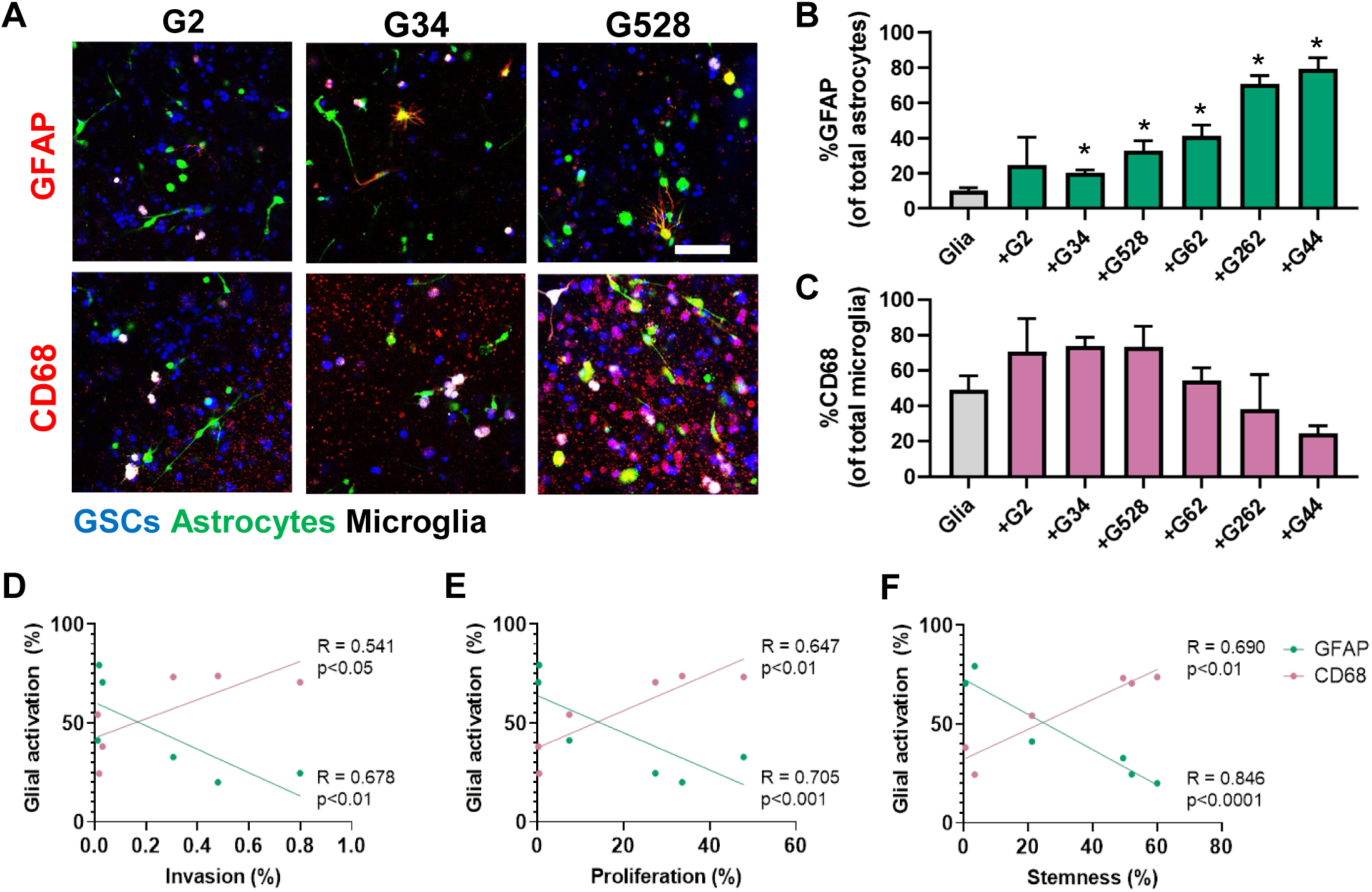
Glial reactivity to glioma cells is patient-dependent, and astrocyte reactivity correlates with glioma CD71 expression. A) Representative immunofluorescence images showing expression of reactivity markers GFAP (red, astrocytes) or CD68 (red, microglia) for glial cells alone (AM) or in tri-culture with G2, G34, and G528. Glioma cells (blue) are labeled only with DAPI, while CellTrackers label the astrocytes (green) and microglia (white). Scale bar is 100 μm. B) Quantified number of GFAP^+^ astrocytes as a percent of total astrocytes. C) Quantified number of CD68^+^ microglia as a percent of total microglia. D) Correlation plot of GFAP vs. CD68 expression. E) Correlation plot of GFAP expression vs. previous GSC Ki67 expression. F) Correlation plot of GFAP expression vs. previous GSC CD71 expression. Pearson’s correlation coefficients and p values are shown in respective plots, with n=3 trials per cell line.

We then performed regression analyses to determine if glial activation correlates with the glioma metrics previously influenced by the cellular TME. Invasion shows a moderate, significant correlation with %GFAP^+^ (R=0.678, p<0.01) and %CD68+ (R=0.541, p<0.05) cells (**Figure 5D**). Proliferation significantly correlates with both: negatively with GFAP (R=0.705, p<0.001) and positively with CD68 (R=0.647, p<0.01) (**Figure 5E**). Because proliferation and stemness positively correlated before, stemness also correlates in similar ways with GFAP+ (R=0.846, p<0.0001) and CD68+ (R=0.690, p<0.01) (**Figure 5F**). Thus, the activation state of the glial cells correlates with all outcomes seen in our systems when analyzed by GSC line.

### In vitro survival does not predict xenograft drug response

Dose response studies are commonly conducted using *in vitro* models that do not effectively recreate the tissue environment or drug delivery method. Our model incorporates interstitial fluid flow to recreate a physiologically-relevant force tightly linked to drug delivery and distribution within the microenvironment. We tested the utility of our system for assessing tumor drug response using a selected panel of clinically relevant therapeutics, including the first-line treatment temozolomide and the following second-line therapies (either in GBM treatment or other central nervous system tumors): carboplatin, methotrexate, etoposide, irinotecan, and BCNU (carmustine). In examining cell survival for both G34 and G528, we found the addition of the TME significantly increased the 50% inhibitory concentration (IC_50_) – with the cells less responsive to almost every therapy when compared to cells in gel alone or in static spheroid culture (no interstitial flow) (**Figure S6**). In several cases, with G34, an IC_50_ could not be calculated due to poor responsiveness in the TME model.

To determine if *in vitro* drug response predicts survival in mice, we implanted either GSC line G34 or G528 into NOD-SCID mice and treated animals with the same therapeutics according to informed doses and schedules shown in **Table S3** (*25–30*). Fewer select drugs were used for G528, and G528-bearing mice generally have longer survival times than those with G34. Temozolomide and BCNU prolonged survival the most (**Figure 6A**), although neither of these therapies had the lowest IC_50_ in any of the models we tested. We then built a proportional hazards model to assess the relationship between average mouse survival time and average *in vitro* survival data at the dose below IC_50_ (of spheroid culture since some IC_50_’s are not determined in the TME model). Based on this hazard ratio model, we find *in vitro* survival data do not predict survival of xenografted mice (**Figure 6B**).

**Figure 6.**
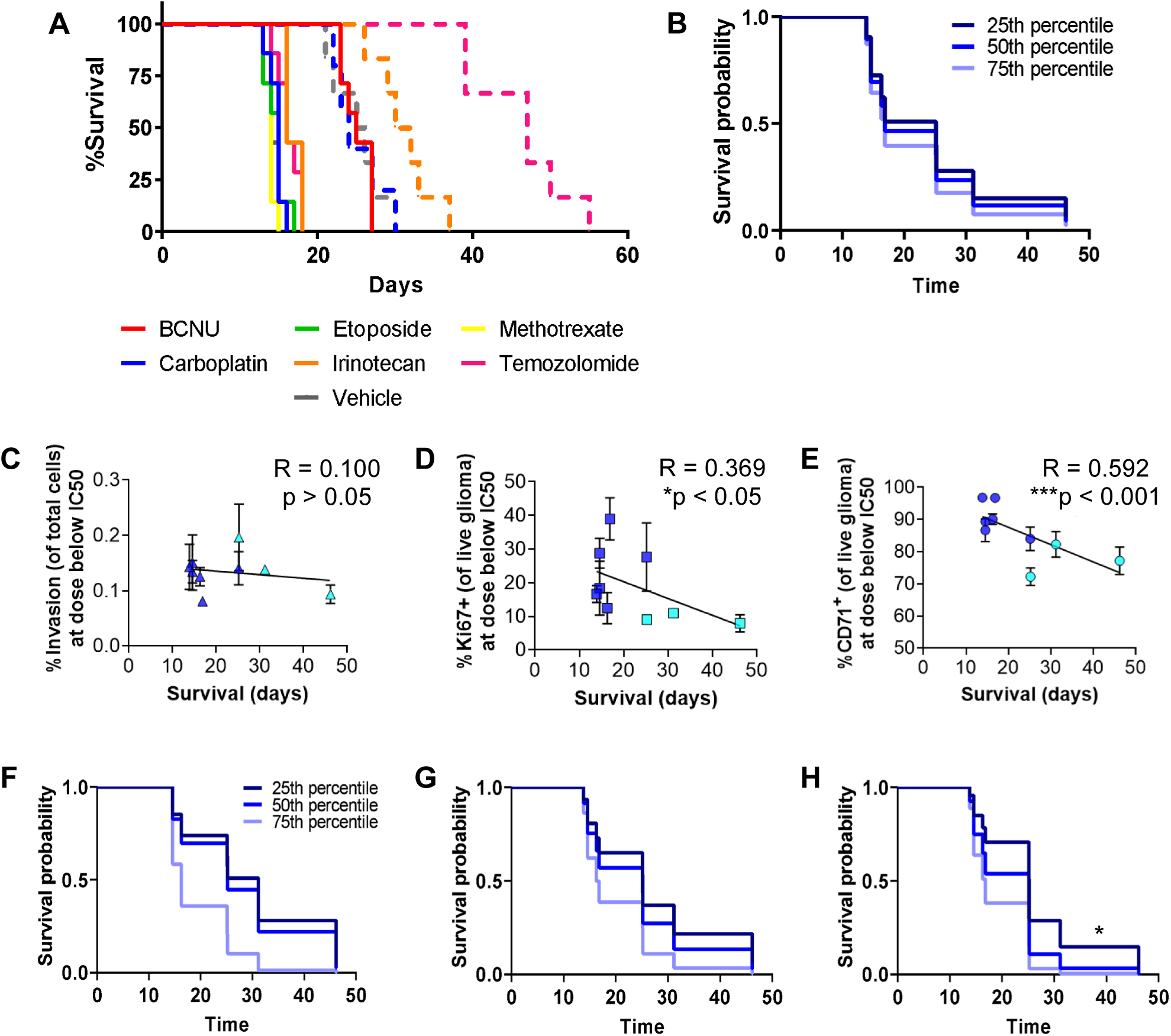
Metrics for proliferation and stemness correlate with xenograft survival, with *in vitro* stemness predicting *in vivo* drug response. A) Kaplan-Meier plots for orthotopic xenografts of 10,000 G34 (solid lines) and 400,000 G528 (dashed lines) treated with chemotherapies at concentrations and regimens based on published literature. G528 was only tested with temozolomide, carboplatin, and irinotecan. B) Survival probability model built using *in vivo* survival data and *in vitro* survival data in the TME model at the dose-below-spheroid-IC_50_ value. C-E) Collective *in vitro* responses at the dose-below-spheroid-IC_50_ value of G34 (dark blue) and G528 (light blue) treated with drugs in the TME model for invasion (C), proliferation (D), and stemness (E). Each data point is n=4, with 6 drug responses for G34 and 3 drug responses for G528. Pearson’s correlation coefficients and p values are shown on respective plots (N=9). F-H) Survival probability models built using *in vivo* survival data and data from (C-E), examining the ability to predict xenograft survival given the percent invasion (F), proliferation (G), and stemness (H). Only stemness was able to predict *in vivo* therapeutic response (by proportional hazards model with *p<0.05).

### Additional malignancy outcomes are necessary to predict drug efficacy

Given the inability of in vitro dose response to predict xenograft survival, we next examined the ability of the other outcomes (invasion, proliferation, and stemness) to predict *in vivo* drug response. We next examined the ability of the other outcomes to predict mouse survival. We again used the values of invasion, proliferation, and stemness at the dose below IC_50_ (from spheroids) since these metrics can decrease as a side effect of decreasing tumor cell number. (*In vitro* data for statistical model development are shown in **Figure S6C-E**.) Regression analysis shows invasion does not correlate with *in vivo* survival (R=0.100; p>0.5) (**Figure 6C**), while both proliferation and stemness significantly negatively correlate with xenograft survival (R=0.369; p<0.05) (**Figure 6D-E;** R=0.369; p<0.05and R=0.592; p<0.001, respectively). A proportional hazards models shows neither invasion (**Figure 6F**) nor proliferation (**Figure 6G**) predict outcomes in mice, but the percentage of CD71^+^ cells does in fact predict *in vivo* mouse survival (p<0.05) (**Figure 6H**). Therefore, glioma stemness in response to drug treatment in our TME model both correlates with and predicts xenograft drug response for two distinct patient-derived cell lines.

## Discussion

The clinical process of characterizing GBM malignancy uses a host of molecular markers, including IDH1-mutant status, MGMT methylation status, and molecular subtypes. While this profiling provides useful information, the resolution is currently not sufficient to accurately and reproducibility predict therapeutic outcomes, necessitating the identification of more prognostic factors and potential predictors of therapeutic response. Developing new and better *in vitro* cancer models will be an important step to identify effective new markers, therapies, and potential means to inform patient-specific drug regimens. In this regard, there is growing appreciation for the role of the tumor microenvironment (TME), both cells and biophysical forces, in cancer progression and therapeutic drug response. Our TME model of human GBM is therefore an advancement toward complementing current molecular profiling using live patient cell responses to microenvironmental factors, and we used this model to study and dissect the discrete effects of glial cells and interstitial fluid flow to glioma progression and therapy.

We analyzed four metrics of glioma behavior in our model, namely invasion, proliferation, stemness, and cell death. These four metrics are often evaluated separately, but proliferation, stemness, and invasion may all be related (*22*). While stemness in cancer is widely debated, *in silico* modeling reveals the need to effectively kill stem-like cells in breast cancer to prevent recurrence (*23*). Furthermore, recent research shows glioma cells do exhibit *de facto* qualities of stemness, including the ability to differentiate into multiple functionally distinct phenotypes recreating a neurodevelopmental hierarchy (*31*). Our patient-derived glioblastoma stem cells were isolated using non-adherent culture, and expression of common stemness markers like CD133, often used for GSC isolation, is not guaranteed in the absence of this positive selection step. Further, expression of the purported stem cell marker CD44 is highly expressed in mesenchymal subtype GSCs but it expressed less in other subtypes, devaluing its use as a general stemness marker. Ultimately, we chose the putative stemness marker CD71 (transferrin receptor) as it is reportedly necessary for GSC maintenance (*32*). We find interstitial fluid flow to have the most consistent effect on increasing stemness as well as proliferation of GSCs.

The cytokine microenvironment and niches that develop within the tumor and brain can be indicative of malignancy and perhaps therapeutic response. We identified increases in CXCL1, CXCL8, and CCL2 upon combining glioma cells with glial cells in our tri-culture model. Importantly, it appears glial cells are the primary source of CCL2, and supplementing GSCs with CCL2 alone reproduced the effects of the TME on glioma invasion and stemness. CCL2 has been shown to be essential for recruiting regulatory immune cells into GBM tumors, and its presence can have negative implications on antiangiogenics and immunotherapies (*33,34*). Thus, the ability to recapitulate a cytokine milieu representative of the *in vivo* tissue environment is uniquely useful for studying anti-cancer therapies and may yield novel insight for the development of combination therapies that are either universal or patient-specific.

Both astrocytes and microglia are known to have pro-tumor phenotypes in glioma, and so an interesting finding within our model is the effects of glioma cells on glial activation. Pan-reactive markers for both astrocytes and microglia changed based on the patient GSC line. Interestingly, astrocyte activation negatively but strongly correlates with microglia activation as well as glioma invasion, proliferation, and stemness, suggesting decreased astrocyte activation is important in glioma progression. We note the activation response tends to cluster by GSC subtype (mesenchymal G2, G34, G62; classical G528; and proneural G262, G44). Similar observations have been made before, as the mesenchymal GBM subtype showed strongest activation of monocytic cells through extracellular vesicles (*35*). A larger set of patient samples will reveal if our model enables similar recapitulation of subtype-dependent interactions.

Towards the goal of therapeutic validation and discovery, we compared our model to previous models for examining chemotherapeutic drug response. The IC_50_ is the concentration necessary to reduce the biological process by 50%, and it can theoretically be applied to any metric. An IC_50_ calculation requires the drug to be able to induce a near-complete response (e.g., 0% cell survival), which we often found difficult in our model due to increased drug resistance in the TME. Furthermore, IC_50_ calculations often did not apply to metrics of invasion, proliferation, and stemness, as these did not consistently decrease toward zero (and sometimes increased) even at high drug concentrations. There are also other metrics – like EC_50_, GI_50_, and GR_50_ – which have different calculation requirements and provide different insight (*36*), suggesting it is worth exploring the potential of these alternative metrics in future analyses.

*In vitro* survival data are often used to justify advancement of drugs to pre-clinical testing, yet we found no potential for *in vitro* survival data to predict *in vivo* xenograft survival across any model tested, including our own (based on %live at dose below IC_50_). Nonetheless, there are several reasons why our survival data may have failed to correlate with *in vivo* results. Our model does not recreate the blood-brain/tumor-barrier, which can impact drug transport and provides different signals to the GBM cells. Our drug dosing concentrations and schedules, while based on prior literature, also may not capture the optimal drug responses. The xenograft model is necessarily in immunocompromised mice and therefore lacks any interactions of the tumor with a functioning immune system. Furthermore, it may not be necessary or desirable to predict xenograft survival since these models can poorly translate to human patients (*12,37*). More work is necessary to understand how best to predict outcomes in patients, and our data suggest examining putative markers of cancer stemness across the platforms may provide important insight toward this aim.

There is a clear need to develop better cancer models, identify new metrics for better response prediction, and improve the resolution and understanding of cancer molecular profiles. Our GBM TME model may help further this goal by enabling recapitulation and study of TME influences on malignancy and drug responses of patient-derived glioma cells. Many models of GBM have been created to study the role of factors like the extracellular matrix and stromal cells (*38–40*), but to our knowledge this is the first model designed for simultaneously probing the effects of three TME components (two glial cell types and interstitial flow) and analysis by multiparametric flow cytometry. Analysis in our model only takes a few days, meaning the largest hurdle is the time required to generate patient-derived cell lines in the clinic. Ultimately, we used this model to identify a potential new metric for evaluating drug responses *in vitro* based on glioma stemness (% CD71^+^), which both correlates with and predicts *in vivo* drug response.

## Materials and Methods

### Study Design

The foundational advancement of this study is using quantification of patient samples for design and development of an engineered tissue model. Glioblastoma resection samples were used to develop a model of the infiltrative brain tumor microenvironment incorporating patient-derived cells, multiple elements of the tumor microenvironment, and cellular ratios representative of actual human patients. Neuropathologists and clinicians were heavily involved in the study design process, including selecting tissue samples with appropriate infiltrative areas, identifying relevant tissue areas for quantification, providing patient cells, selecting a panel of clinical drugs, and informing metric evaluation. The hydrogel material used here was based on previous studies wherein the composition was optimized to achieve relevant rates of interstitial fluid flow (*8*).

### Patient immunohistochemistry and image analysis

Patient samples are accessed through the University of Virginia Biorepository and Tissue Research Facility and selected by a neuropathologist (J. Mandell) based on a definitive diagnosis of glioblastoma (World Health Organization grade IV). All patients had completed tumor resections at the University of Virginia between 2010 and 2013. Samples were de-identified and processed to select tumor sections that included a portion of adjacent non-bulk tumor tissue (here referred to as the parenchyma interface) as identified by a neuropathologist (F. Bafakih)(*20*).

Formalin fixed paraffin embedded 8μm sections are deparaffinized with xylene and rehydrated in graded ethanol, antigen retrieved using high pH antigen unmasking solution (Vector Labs), and stained with anti-ALDH1L1 (Abcam) and anti-Iba1 (Abcam), followed by DAB substrate (Vector) according to manufacturer suggested protocols and counterstained with hematoxylin (Thermo Scientific). Hematoxylin and eosin staining was performed by the University of Virginia Biorepository and Tissue Research Facility following standard protocols. Areas at the tumor-parenchyma invasive front of tumor resections are imaged using wide-field microscopy with EVOS FL Auto (Life technologies) and Aperio Scanscope (Leica Biosystems) and quantified using ImageJ (National Institutes of Health). Cell populations are reported as a percentage of total cells identified by the nuclear counterstain.

### Cell lines and culture

Patient-derived human glioblastoma stem cells (GSCs) were a generous gift to Benjamin Purow from Jakub Godlewski and Ichiro Nakano (who derived them while at Ohio State University). These cells (G2, G34, G44, G62, G262, G267, and G528) are maintained in non-treated culture flasks in Neurobasal medium (Life Technologies) supplemented with 1% B27, 0.5% N2, 0.01% FGF, 0.1% EGF, 0.3% L-Glutamine, and 1% penicillin-streptomycin. Human primary cortical astrocytes are purchased from Sciencell and cultured according to manufacturer’s suggested protocol. Human SV40-immortalized microglia are purchased from Applied Biological Materials, Inc and cultured in Dulbecco’s Modified Eagle’s Medium (DMEM; Life technologies) supplemented with 10% fetal bovine serum (FBS). All cell lines are maintained at 37°C in a humidified incubator containing 5% CO_2_ and 21% O_2_ and tested annually for mycoplasma (negative).

### Three-dimensional cell assays

Experiments are carried out with 8μm pore size tissue culture inserts (Sigma Aldrich). Cells are fluorescently labeled with CellTracker dyes (Life technologies) and Vybrant dyes (Life technologies) according to manufacturer suggested protocol. Glioblastoma cells (5.0×10^5^), astrocytes (8.0×10^4^), and microglia (8.0×10^4^) are seeded in 75 μL gel comprising (0.2% hyaluronan; ESI Bio) and 0.12% rat tail collagen I (Corning) using cell ratios quantified from human sections. Gels solidified at 37°C in a humidified incubator containing 5% CO_2_ and 21% O_2_ for 3 hr, then serum-free medium (Astrocyte Basal; Sciencell, with 1% B27, 0.5% N2) was added to the top and bottom of each tissue culture insert such that medium level was consistent inside and outside the insert.

### In vitro drug dosing experiments

For screening studies, 24hrs after gels are seeded into transwells, a range of concentrations of BCNU, carboplatin, etoposide, methotrexate, irinotecan, and temozolomide chemotherapies are added on top of the gels (pressure head of 1 cm) to drive flow through the gels, leading to an average velocity of 0.7 μm/s. A small volume (25 μL) of chemotherapeutic-free media was added to the bottom compartment. After 24hours of dosing, media that had flowed through the gel into the bottom compartment was carefully removed, and the same range of concentrations of each drug was added again at the top to reestablish the pressure head for another 18hrs of dosing. The cells are then collected for flow cytometry and the membranes fixed for invasion analysis.

### Invasion assay and flow cytometry

After 18hr, gels are removed from tissue culture inserts and digested using Roche Liberase DL (Sigma Aldrich). Cells migrating through the porous membrane are identified by staining with 4′,6-diamidino-2-phenylindole (DAPI; Invitrogen), counting five representative fields per insert, and reported as total cells invaded/total cells seeded x 100 (%) for each insert. Cells remaining post-gel digestion are stained for Live/dead (Life technologies), CD71 (eBioscience), and Ki-67 (eBioscience) according to manufacturer’s suggested protocol. Flow cytometry was performed using Guava easyCyte 8HT (Millipore) and analyzed using guavaSoft 2.7 (Millipore).

### Tumor inoculation in animal studies

All animal procedures are conducted in accordance with the University of Virginia Institutional Animal Care and Use Committee (Charlottesville, VA). 8-10week old male NOD-SCID mice are inoculated with 10,000 GSCs derived from patient G34 (n=7) or 400,000 GSCs derived from patient G528 (n=6) resuspended in 10μL of neurobasal media supplemented with N2, B27 without vitamin A, and glutamax. Inoculations are performed 2mm lateral and posterior to bregma at a depth of 2.2mm. Seven days after inoculation, chemotherapeutics are administered intraperitoneally according to Table S3. Animals are assessed daily for signs of distress and are euthanized accordingly when they displayed humane endpoint criteria.

### Proportional hazards model development

Data used in this analysis includes 2 GSC lines (G34 and G528) and 6 treatments (BCNU, Carboplatin, Etoposide, Irinotecan, Methotrexate, and Temozolomide). Measures of viability, invasion, proliferation, and stemness are measured (n=4 biological replicates with 3 technical replicates each). Survival of mice with these cell lines and treatments was assessed, where mice exposed to the G34 cell line are treated with all 6 treatments (n=7 mice with each treatment type) and mice exposed to the G528 cell line are treated with Carboplatin (n=5 mice), Irinotecan (n=6 mice), and Temozolomide (n=6 mice). To assess a relationship between mouse survival and measures derived from experiments (viability, invasion, proliferation, and stemness), averages across all replicates (within cell line and treatment type) from experiments and across mice are calculated for modeling. Proportional hazards models assess the relationship between average mouse survival time and average experiment outcomes. Due to the sample size, only univariate models are considered. The hazard ratios presented indicate the change risk of death for a change of 10% of the range of variable considered. For example, viability measurements ranged from 67.44 to 96.85, a range of 29.41 units. Figures display the model predicted survival curves for a patient with low, medium, or high values for the outcomes of interest. Low, medium, and high are defined by the 25th, 50th, and 75th percentiles of the data.

### Immunostaining of tissue samples

Tissue samples are collected, soaked in sucrose, cryoembedded, and sectioned at 12μm using a Leica CM 1950. Three sections at varying depths within the tumor are immunostained with mouse anti-human nuclei (clone 235-1, Millipore) followed by secondary Dylight 488 horse anti-mouse (Vector), rat Ki67 conjugated to eFluor570 (SolA15, eBioscience), and rabbit Sox2 (Millipore) followed by secondary AlexaFluor 660 goat anti-rabbit (Life technologies).

### Design of experiments analysis

JMP software (SAS) was used to identify key differences among experimental conditions. Independent variables included patient from which the glioblastoma stem cell is derived from (GSC), addition of each of the glial cell microenvironmental conditions (cells), and interstitial flow (flow). Dependent variables are outcome measures of invasion, proliferation, and stemness. The classical screening design was fit for standard least squares to determine which factors have the main effect, and the resulting effects are summarized. A value of p<0.05 was considered statistically significant.

### Statistical modeling of experimental outcomes

Quantile regression of the median was used to assess the relationship between experimental conditions and outcomes of interest. Experimental conditions include glioblastoma stem cell line, glial cell, and interstitial flow. Outcomes include invasion, proliferation, and stemness. All Interactions between experimental conditions were included in the models. Quantile regression analysis was performed using the QUANTREG procedure in SAS 9.4 (Cary, NC). A value of p<0.05 was considered statistically significant.

### Statistical analysis

All *in vitro* results are repeated at least three times, and at least five animals are used for *in vivo* results to yield sufficient biological replicates based on power analyses. All data are presented as mean ± standard error of the mean (SEM). Independent, paired t tests are used to compare all *in vitro* results, with analyses for invasion and cell death conducted as ratio-paired tests. Independent, unpaired t tests and two-way ANOVA was used for statistical analysis of unmatched groups (in vitro glial activation and in vivo analyses). All dot plots, including Kaplan Meyer curves, and statistical analyses are generated or performed using Graphpad Prism software, respectively. A value of p<0.05 was considered statistically significant. Statistically significant differences are determined by ANOVA followed by Tukey’s t tests.

## H2: Supplementary Materials and Methods

### CellTracker and media viability optimization

Cells are fluorescently labeled with a range of concentrations of various CellTracker dyes (Life technologies) and Vybrant dyes (Life technologies) according to manufacturer’s suggested protocol and maintained in respective serum-free media. Growth of labeled cells was measured after 18 hours, 48 hours, and 72 hours using the Cell Counting Kit-8 (CCK-8) cell proliferation and cytotoxicity (Dojindo) according to manufacturer’s suggested protocol. After 72 hr, cells are also assessed for viability using Live and Dead ReadyProbes Reagents (Life technologies) and imaged using wide-field microscopy with EVOS FL Auto (Life technologies) and quantified using ImageJ (National Institutes of Health). Each CellTracker or VybrantDye test performed similarly for each glial cell type (Figure S2). Cells are also tested in varying media compositions to determine optimal viability using the previously described assays. Media compositions tested include basal astrocyte medium (Sciencell), supplemented with 1% B27 and 0.5% N2, and/or 0.01% FGF and 0.1% EGF.

### Optimizing parameters that affect tumor cell viability in tri-culture

We used our histological cellular analysis to develop a model of the human GBM tumor microenvironment incorporating patient-derived glioma stem cells (GSCs), astrocytes, and microglia. The cells are mixed at a ratio of 6:1:1, respectively, with 1 million GSCs/mL. In this model, we first examined how 7 different patient-derived GSCs responded to the cellular tumor microenvironment. **Table S1** shows a summary of patient cell properties. We first examined glioma cell viability in tri-culture using CCK-8 assays for three different media formulations based on Astrocyte Basal Medium (**Figure S2**). For G34 cells, astrocyte basal medium and medium supplemented with EGF and bFGF, two growth factors used in culture and maintenance of the GSCs, performed the worst with a starting GSC viability of less than 60% on day one, and further decreasing viability with additional days.

Supplementing the growth factors with B27 and N2, common culture reagents for GSCs and neuronal cells, showed similar starting viability with improved maintenance over time. The best medium formulation was Astrocyte Basal medium supplemented with only B27 and N2, which maintained glioma cell viability around 80% for up to 3 days of tri-culture. We similarly optimized re-harvesting of the cells from the hyaluronan gels for subsequent staining and assessment by flow cytometry. Of three enzyme formulations tested, Liberase DL maintained glioma cell viability the best and was used for all further gel degradations (**Figure S2**).

### Sex profiling of patient cells

Cells are incubated with 50 ng/mL colcemid (Karyomax; Invitrogen) 4-6 hr prior to fixation to enrich for mitotic cells. The cells are collected and centrifuged at 1000 rpm for 5 min (used for all subsequent centrifugation steps). The cell pellet was washed once with PBS and centrifuged again. The cells are resuspended in 0.075 M KCl and incubated at 37°C for 18 min; then 0.5 mL of freshly prepared fixative (3:1 methanol-glacial acetic acid) was added before centrifugation. The cells are resuspended in fixative added drop-wise and incubated at room temperature for 15 mins before centrifugation. Cells are suspended in a final volume of 0.3-6 mL fixative (added drop-wise; final volume based on pellet size) and 12 μL is dropped onto microscope slides, which are then air-dried overnight. Human X/Y centromere enumeration probes (Metasystems Probes) are added to the sample, sealed under a coverslip with rubber cement, and placed on a hotplate at 75°C for 3 mins for probe and sample denaturation. Samples are placed in a humidified incubator at 37°C for 4-6 hr to allow probe hybridization. After removing the coverslip and any glue remnants, samples are washed in 0.4X SSC (pH 7.0) at 72°C for 2 mins and 2X SSC, 0.05% Tween-20 (pH 7.0) at room temperature for 30 seconds. The slides are rinsed briefly in distilled water and allowed to air dry. Antifade solution (90% glycerol and 0.5% N-propyl gallate) with 300 nM DAPI was added to the slides, sealed under a 22×50 mm coverslip (Corning Incorporated) with nail polish, and incubated at room temperature for 10 mins prior to analysis on a Nikon Eclipse Ti inverted microscope (Nikon Instruments Inc., NY, USA) equipped with ProScan automated stage (Prior Scientific), Lumen200PRO light source (Prior Scientific), CoolSNAP HQ2 CCD camera (Photometrics), and a 60X/1.4 NA Plan-Apochromatic objective.

## General

The authors would like to thank Nikhith Kalkunte and the flow cytometry core and Biorepository and Tissue Research Facility at the University of Virginia for their assistance with data collection.

## Author contributions

JMM, JXY, and RCC conceived the experimental design; RCC and JXY conducted the experiments; RCC, JXY, KMT, AP, GFB, KMK, MB, SCS, ALB, JWM, and BH collected or analyzed the data; JWM BWP, FFB, DC, BH, and JMM provided resources and guidance on data collection and interpretation; RCC, JXY, and JMM wrote the manuscript; and all authors provided critical manuscript editing and feedback.

## Supplementary Figures

**Figure S1.**
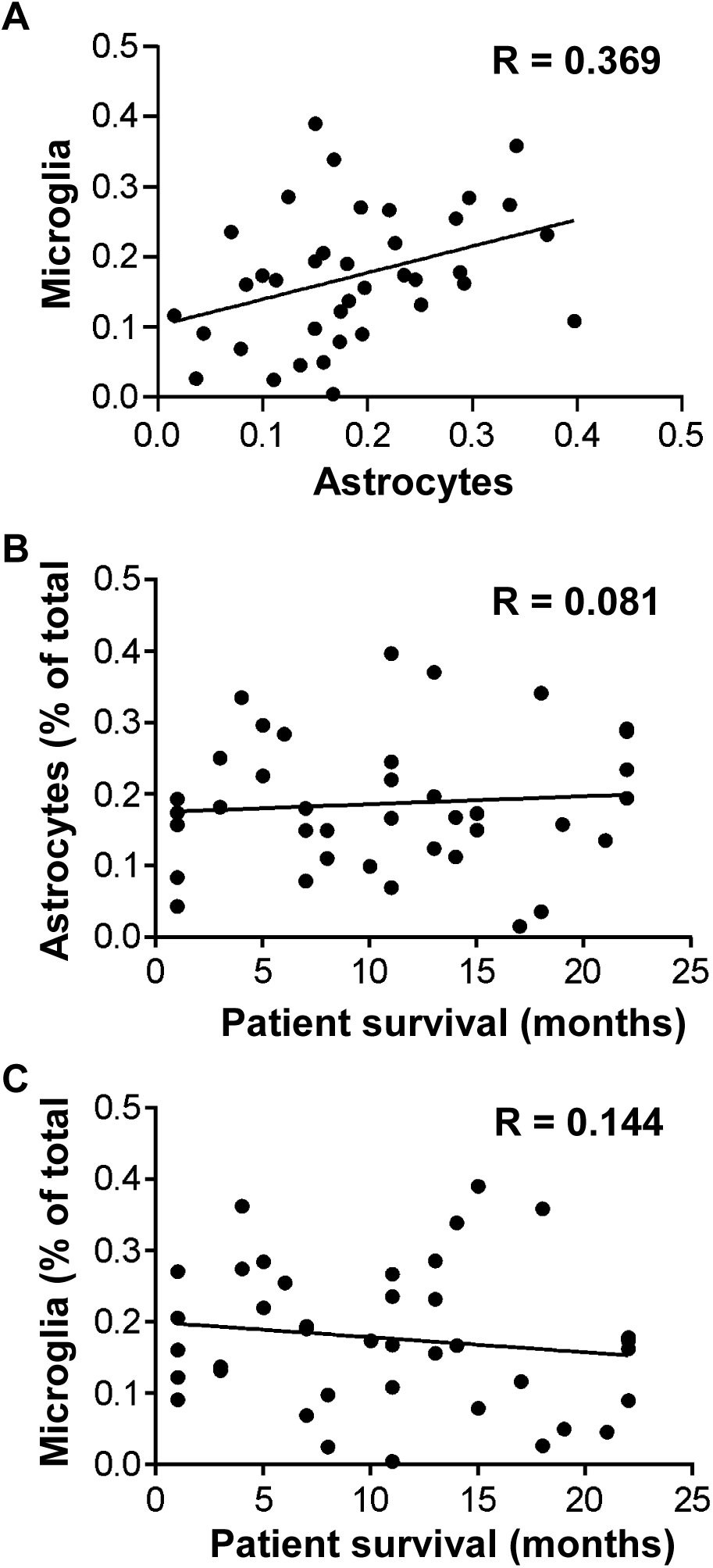
Correlations of glial cell number fraction with each other and overall patient survival. A) Correlation plot showing numbers of microglia (Iba1^+^) versus astrocytes (ALDH1L1^+^) for each analyzed patient sample. B) Correlation plot of astrocyte number versus respective patient survival. C) Correlation plot of microglia number versus respective patient survival.

**Figure S2.**
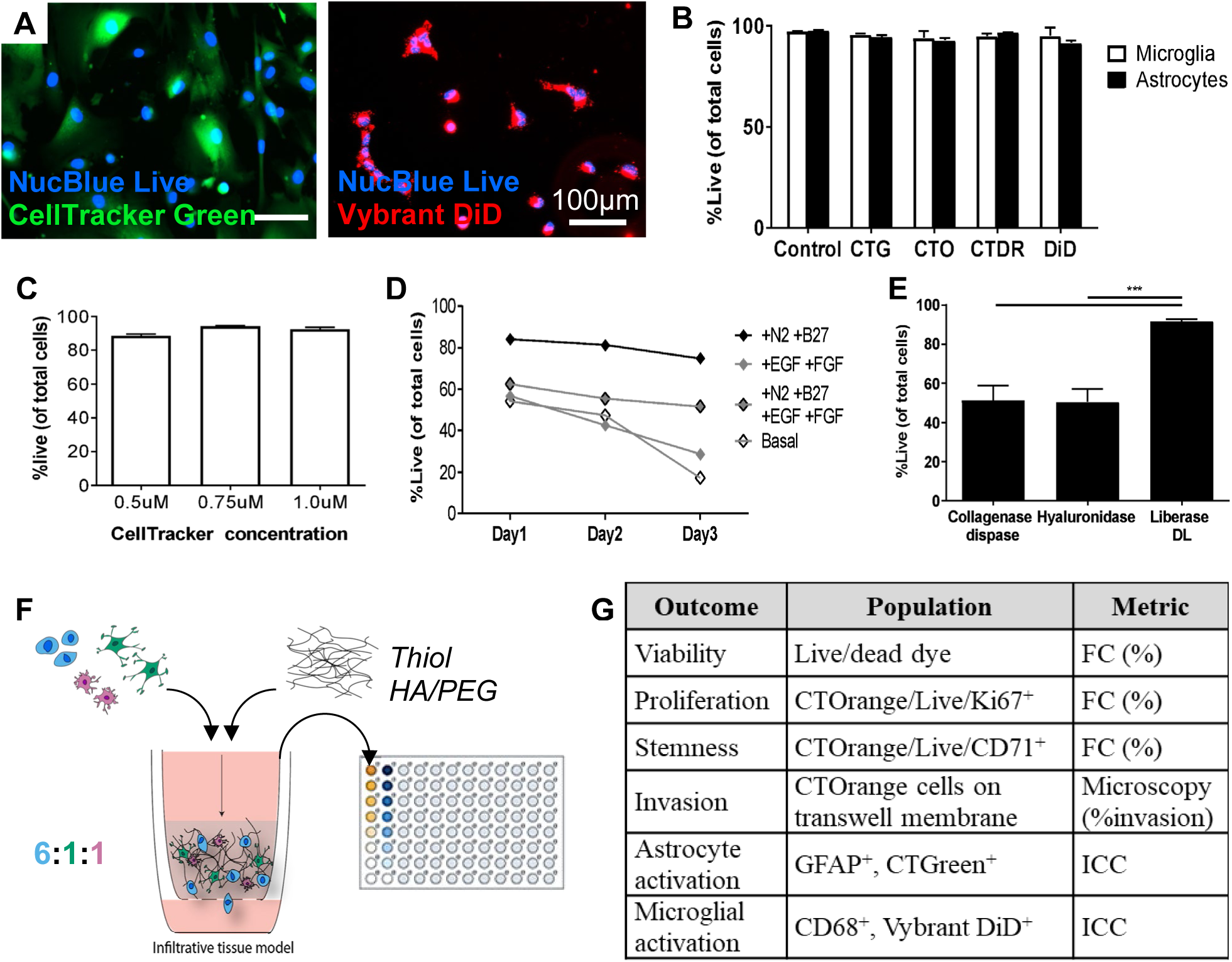
Optimization of the infiltrative human glioblastoma TME model. A) Representative fluorescence images of human astrocytes labeled with CellTracker Green (left) and human microglia labeled with Vybrant DiD (right). B) Glial cell survival following labeling with various cell dyes. C) Glial cell survival with different concetrations of CellTracker dyes. D) Glioma cell survival in tri-culture under different media conditions. E) Glioma cell survival after digestion and cell pelleting of tri-culture hydrogels using different enzyme treatments. F) Schematic showing elemental construction of the infiltrative TME model, where glioma cells are blue, astrocytes are green, and microglia are pink. G) Summary of outcomes and metrics used to evaluate the infiltrative TME model.

**Figure S3.**
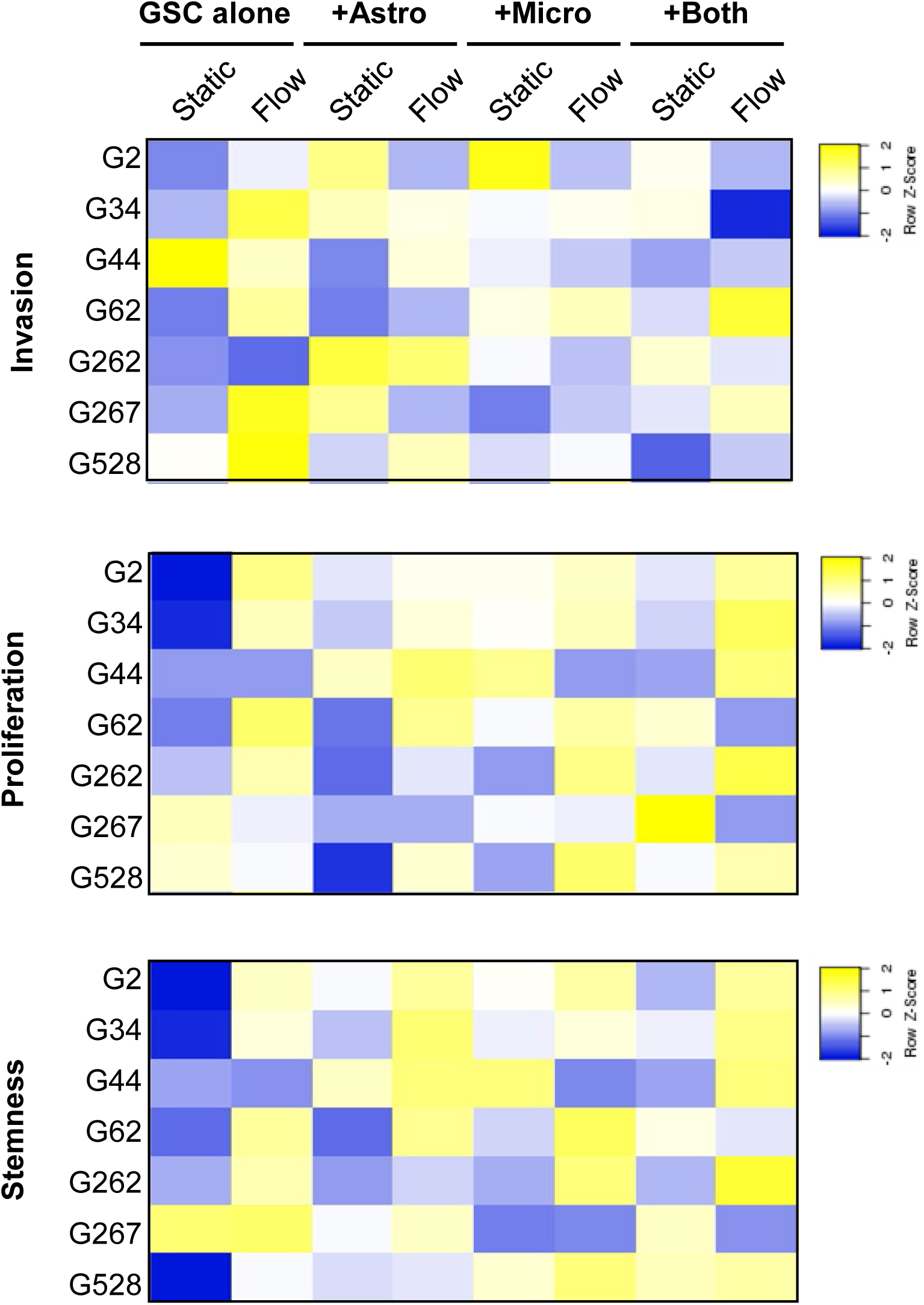
Heatmap representations of changes in glioma cell metrics (invasion, proliferation, and stemness) in response to each combination of TME elements. Data are represented on a normalized z-score from −2 (blue) to +2 (yellow).

**Figure S4.**
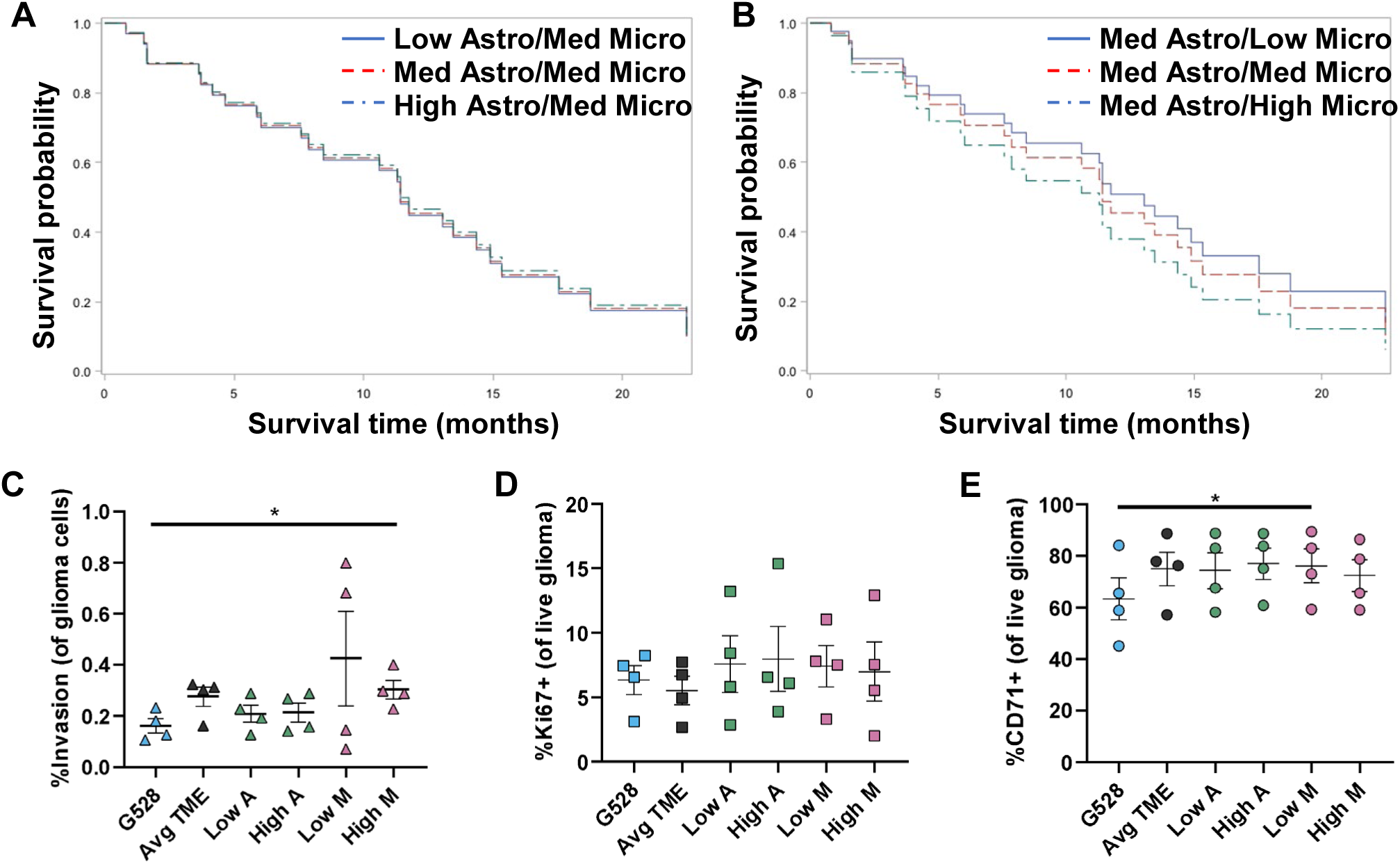
Ratiometric tuning of glial cell numbers influences glioma cell outcomes. Analyzed patient samples were grouped based on the numbers of astrocytes and microglia quantified in infiltrative regions, and proportional hazards models were built to predict patient survival based on TME composition. Resulting Kaplan-Meier curves for varied astrocyte numbers (A) and microglia numbers (B) show microglia have the strongest effect on overall survival. In our TME model, we varied our standard glial ratio (1:1) by 25% to represent high or low ratios (e.g., 0.75:1 for low astrocytes and medium microglia), and examined the effects on glioma cell invasion (C), proliferation (D), and stemness (E).

**Figure S5.**
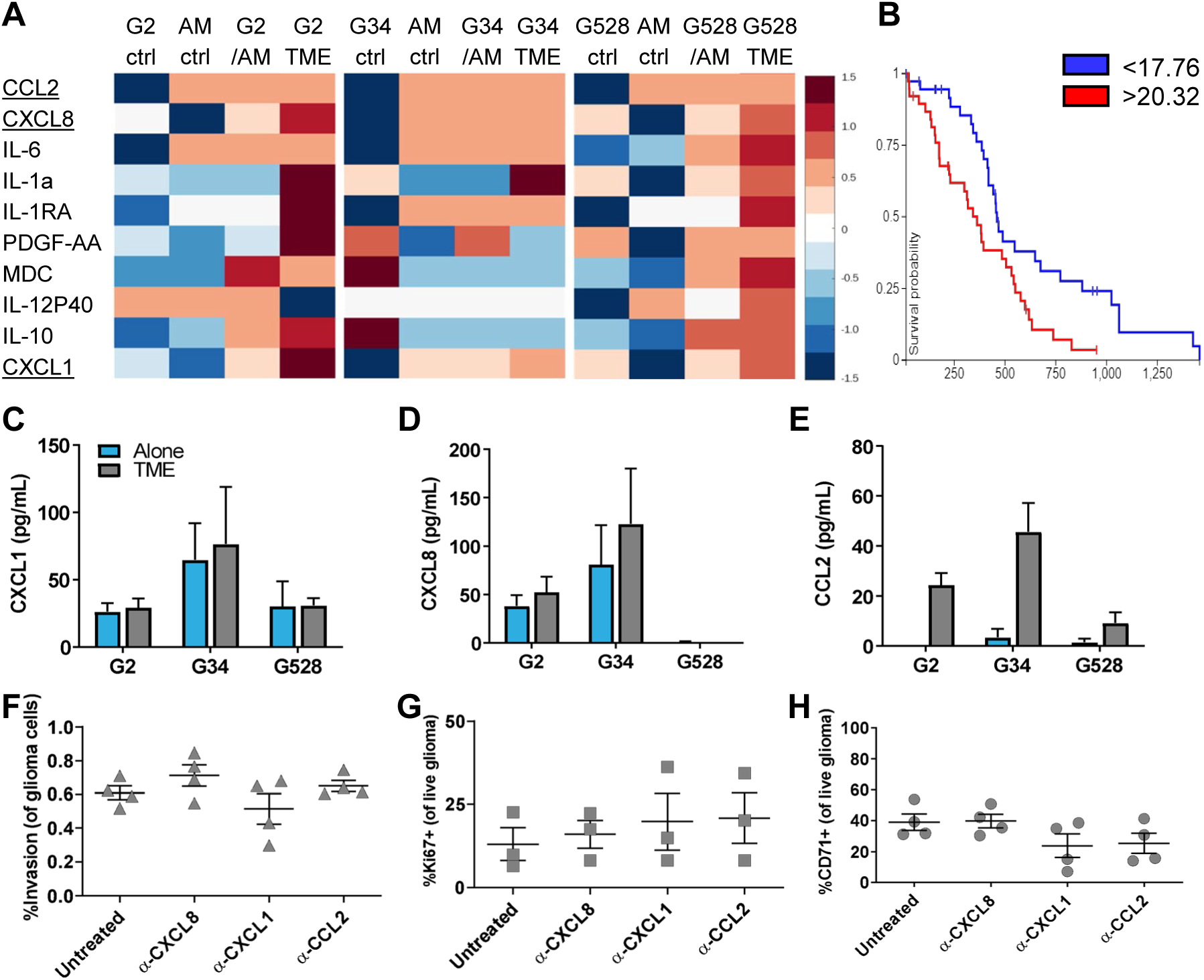
Cytokine contributions from the cellular TME. A) Heatmap of cytokine data obtained using a Luminex array. CCL2 (MCP1), CXCL8 (IL8), and CCL1 (GRO1) were further explored based on high expression in the GSC+TME conditions. B) Survival data from The Cancer Genome Atlas for CCL2, showing survival of patients in the highest and lowest quartiles of CCL2 expression (p<0.01; n=38). C-E) Quantification of cytokines in tri-culture by ELISA for CXCL1 (C), CXCL8 (D), and CCL2 (E). (F-H) Blocking studies performed using antibodies against the identified cytokines and subsequent assessment of glioma cell invasion (F), proliferation (G), and stemness (H). Cytokine blocking showed minimal effect on all metrics in contrast to receptor blocking, possibly due to limitations in antibody transport (vs. cytokines) or concentration effects.

**Figure S6.**
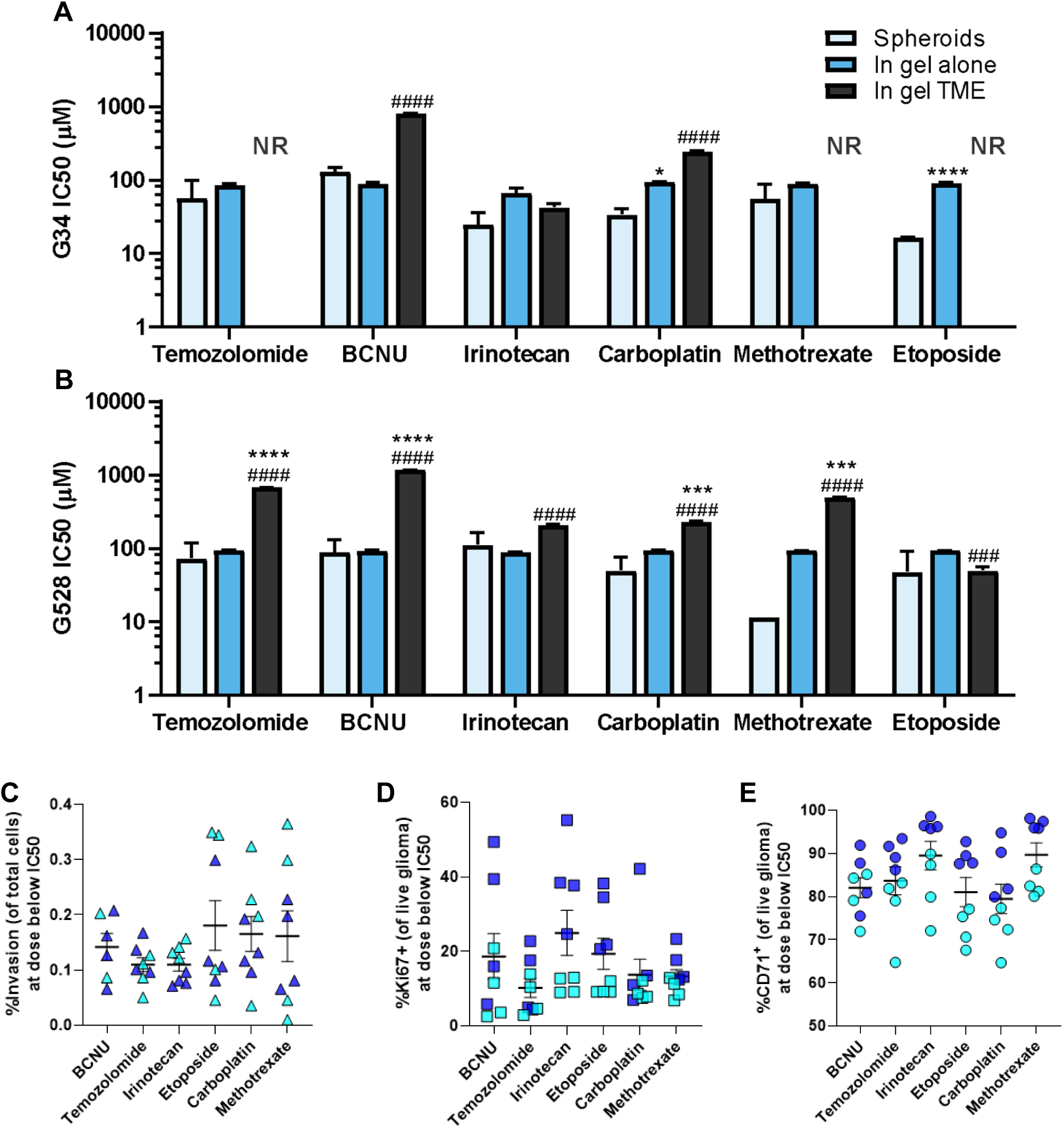
Drug response *in vitro* data. IC_50_ data based on cell survival for six different drugs with G34 (A) and G528 (B) cells cultured either as spheroids, in 3D hydrogels alone, or in 3D hydrogel tri-cultures with glial cells. NR = IC_50_ concentration Not Reached up to 1000 μM. Data are shown on log scale. Statistics conducted using paired t-tests. Asterisks (*) show comparisons to spheroids data and pound symbols (#) show comparisons to ‘in gel alone’ data, with *p<0.05, ***p<0.001, and ****p<0.0001. (C-E) We also assessed glioma cell invasion (C), proliferation (D), and stemness (E) in tri-culture at the drug dose below (spheroid) IC_50_. G34 are shown in light blue, and G528 are shown in dark blue. This data was used to generate the proportional hazards model prediction of xenograft survival.

## Supplementary Tables

**Table S1.**
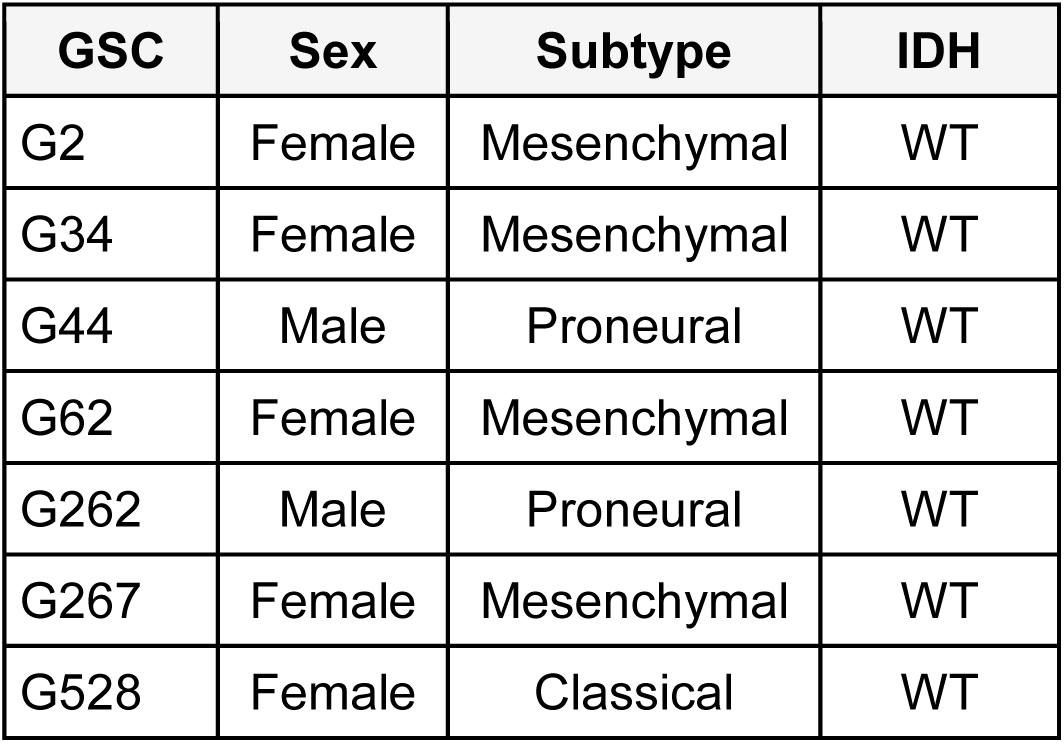
Characteristics of patient-derived glioma stem cells.

**Table S2.**
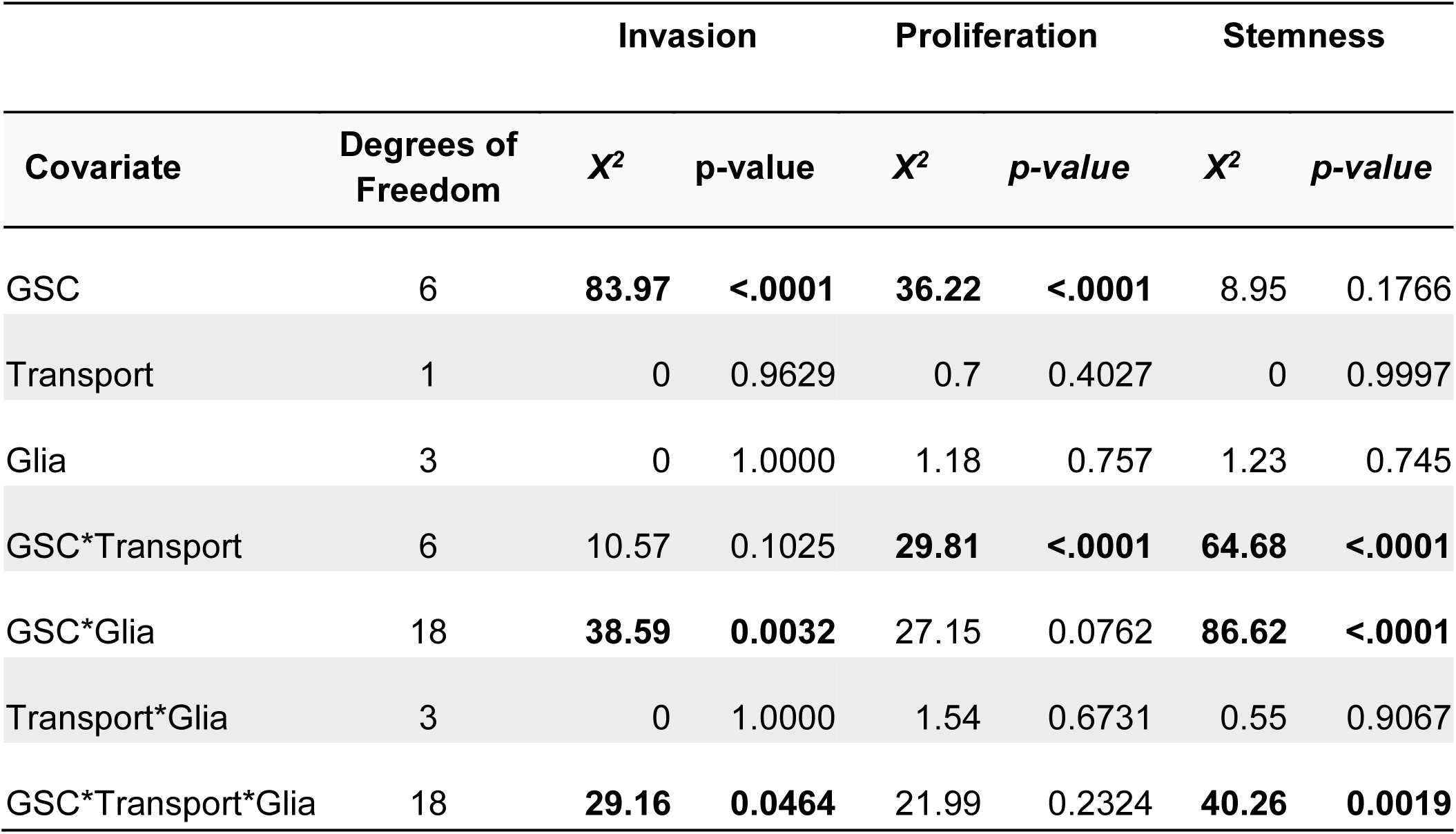
Quantile regression predictive modeling of covariates in the in vitro model.

**Table S3.** *In vivo* drug dosing paradigm.

| Drug | Dose (mg/kg) | Concentration (mg/ml) | Schedule (days administered) | Reference |
| --- | --- | --- | --- | --- |
| Temozolomide | 5 | 1 | 7,8,9,10,11 | 23, 26 |
| BCNU (carmustine) | 25 | 5 | 7, 10 | 24 |
| Carboplatin | 10 | 2 | 7, 10 | 28 |
| Etoposide | 3 | 0.6 | 7,8,9,10,11 | 27 |
| Irinotecan | 4 | 0.8 | 7,8,9,10,11 | 25 |
| Methotrexate | 25 | 5 | 7, 10 | 28 |
| Vehicle (10% DMSO in saline) | 5 | 1 | 7, 10 |  |

